# Engineering protein expression dynamics with Tet-ON and dTAG degron systems: from precise control to oscillations

**DOI:** 10.64898/2025.12.16.694651

**Authors:** Benjamin Noble, Oliver Cottrell, Andrew Rowntree, Veronica Biga, Florence Woods, Xinjie Wang, Nancy Papalopulu, Anzy Miller

## Abstract

Precise temporal control of protein expression is essential for dissecting protein function and dynamic cellular processes. We present a framework for engineering tunable oscillatory protein expression (repeated pulses in expression) using widely adopted molecular tools, applying them to modulate NGN3 expression. Single-cell time-lapse microscopy reveals that the Tet-On system unexpectedly generates asynchronous oscillations in protein expression under continuous doxycycline administration. These oscillations are dependent on protein instability and are not tunable by doxycycline concentration. In contrast, the dTAG degron system enables precise, reversible, concentration-dependent control of protein degradation and reaccumulation. Coupled with a constitutive promoter, we achieve synchronous oscillatory protein expression (COD: **C**onstituitive promoter driving **O**scillations via **D**egradation). Mathematical modelling identifies optimal dTAG drug addition and removal timings using the COD system to flexibly tune NGN3 oscillation periods while maintaining other oscillatory parameters (mean level and peak-to-trough fold-change). Using microfluidics (COD+CHIPS) we reproduce model-predicted 5 and 10 hours periodicities while maintaining similar mean levels and peak-to-trough fold-changes. This work introduces a generalisable, programmable approach for generating and modulating protein oscillations, allowing investigation into how dynamic protein expression governs cellular function.

## INTRODUCTION

The ability to control the expression of proteins is required to effectively interrogate protein function, and, in turn, uncover biological mechanisms. Advances in molecular and synthetic biology have provided powerful tools to manipulate protein abundance, including inducible promoters, knock-outs, knock-downs and targeted degradation systems (reviewed in Eisenhut et al. 2024). However, it is not always possible or it is unclear if these approaches have the ability to be reversible, able to fine-tune protein levels or the ability to control protein levels repeatedly. Therefore, controlling protein expression dynamics has been largely overlooked. This is an important limitation, since proteins are not always steadily expressed, but instead can be highly dynamic. In particular, many proteins show repeated peaks and troughs in protein levels over time (oscillations), from longer 24 hour cycles (circadian) to shorter ultradian oscillators (less than 24 hour periodicity; Maeda and Kageyama 2024; Yagita 2024). These include many key developmental transcription factors, signalling effectors and proteins important in cancer and in response to stresses (Sabherwal et al. 2021; Cottrell et al. 2025; Sonnen and Janda 2021; Purvis and Lahav 2013; Carraco, Martins-Jesus, and Andrade 2022; Isomura and Kageyama 2014). For example, transcription factors and signalling molecules downstream of the NOTCH signalling pathway have been found to oscillate during diverse developmental contexts such as neurogenesis, pancreas development, somitogenesis, myogenesis, embryonic stem cell differentiation and intestinal cell differentiation (Miller et al. 2025; Weterings et al. 2024; Imayoshi et al. 2013; Seymour et al. 2020; Y. Zhang et al. 2021; Kobayashi and Kageyama 2010; Isomura and Kageyama 2025; Maeda and Kageyama 2024; Lahmann et al. 2019). These oscillations have been shown to be important, for example controlling cell fate decisions, the timing of differentiation, balancing proliferation with differentiation, transitioning from progenitor to differentiation states, controlling the spatiotemporal progression of development and stem cell exit from quiescence (Miller et al. 2025; Weterings et al. 2024; Marinopoulou et al. 2021; Imayoshi et al. 2013; Seymour et al. 2020; Shimojo, Ohtsuka, and Kageyama 2008; Y. Zhang et al. 2021; Soto et al. 2020; Hubaud and Pourquié 2014). In addition, the different parameters of oscillations (period, peak-to-trough fold-change, phase) have been shown to be decoded (Goentoro and Kirschner 2009; Dean et al. 2024; Miller et al. 2025; Schröter and Oates 2010; Weterings et al. 2024; Sonnen, Lauschke, et al. 2018; Sabherwal et al. 2021), which highlights a critical need for tools that can generate precisely controlled oscillatory dynamics, in order to test the function of different dynamic regimes.

Very few approaches have been developed to create oscillatory protein expression and functionally interrogate protein dynamics. One optogenetic system uses blue light to activate gene expression with temporal and spatial precision, and has been applied to compare oscillatory and sustained expression, and to create oscillations with differing period lengths (Imayoshi et al. 2013; Isomura, Ogushi, et al. 2017; Shimojo, Isomura, et al. 2016). However, blue light can cause significant toxicity, leading to cellular damage and death (Duke et al. 2020; Wäldchen et al. 2015; Opländer et al. 2011). In addition, the blue light wavelengths used could interfere with researcher’s existing experimental setups, for example exciting fluorescent proteins such as eGFP which are commonly used as reporters (Hadjantonakis and Nagy 2001). Another strategy has employed pulsing of small-molecule inhibitors/activators of signalling pathways or cytokines to entrain endogenous oscillations. Pulsing of modulators of the NOTCH, WNT and ERK pathways (DAPT, Chiron and SU5402 respectively) has allowed modulation of the phase or the period of oscillations (Sonnen, Lauschke, et al. 2018; Weterings et al. 2024; Simsek et al. 2023), and pulsing cytokines has enabled entrainment of oscillations as well as the ability to vary periods and amplitudes (Sorre et al. 2014; Kellogg and Tay 2015). However, these systems are limited by its reliance on the availability of specific commercial inhibitors/agonists/cytokines for the protein of interest. Likewise, circadian biologists are able to synchronise circadian oscillations between cells via glucocorticoid or temperature pulsing (Feillet et al. 2014; Brown et al. 2002), but these are also indirect manipulation methods and likely to cause other systemic effects. Therefore, there remains a need for generalisable tools to achieve precise temporal control over the dynamics of any protein of interest, whereby we are able to create oscillatory protein expression with the added ability to fine-tune parameters of the oscillations (for example; period, amplitude/fold-change, mean level or phase of the oscillations).

We turned to widely used systems for inducible gene expression and inducible degradation, and asked whether their capabilities could be extended to achieve finely tuned and reversible temporal control at the single cell level. We reasoned that, to be able to create oscillatory expression from scratch (of any protein of interest), we needed control over protein production (to create peaks in expression) and protein degradation (to create troughs), along with the capacity to repeatedly alternate between these states across multiple cycles. Therefore, we interrogated two systems, one that regulates the activation of gene expression (Tet-On system) and one that regulates the stability of the protein post-expression (dTAG system), and asked whether each system individually, or combined could create tunable oscillatory expression: in particular, whether oscillations of different period lengths could be achieved.

The Tet-On/Off system is one of the most widely used gene expression systems in mammalian cells (T Das, Tenenbaum, and Berkhout 2016; Gossen and Bujard 2001). In this system, activation of expression of an exogenous protein of interest is controlled by the addition (Tet-On) or removal (Tet-Off) of Tetracyclines or derivatives (e.g. Doxycycline). Over time, several version improvements of the system have been developed, offering reduced background and high induced gene expression levels (T Das, Tenenbaum, and Berkhout 2016). As a result, the Tet-On/Off system has become a staple for inducing gene expression worldwide, which is why we chose it to investigate it here.

The degradation tag (dTAG) system allows a protein of interest, tagged with a short epitope tag (FKBP12 point mutant (FKBP12^F36V^, simplified to FKBP-V), to be specifically and rapidly degraded upon addition of a small molecule dTAG drug (Nabet, Ferguson, et al. 2020; Nabet, Roberts, et al. 2018). The binding of the drug to the tag results in recruitment of an endogenous E3 ligase and ubiquitination of the FKBP-V-tagged protein and subsequent degradation by the proteasome. This system has become widely adopted, effectively degrading proteins across diverse developmental and disease models (Mehta et al. 2023; Nabet 2021), and was chosen in this study for its ability to directly control the degradation of our protein of interest.

Although both dTAG and Tet-On systems are widely used, there is insufficient characterisation related to reversibility and responsiveness at the single cell level. Therefore, here, using time-lapse single-cell microscopy, we show how these systems can be manipulated to achieve precise and temporal control of protein expression such that oscillatory expression of different characteristics can be created at will. For the Tet-On system, we unexpectedly find that constant doxycycline administration results in oscillatory expression in the majority of cells that respond, but is asynchronous across the population, and a feature invisible in bulk population analysis. We find that changing the doxycycline concentration does not provide control over the oscillatory dynamics or level, but instead affects the number of cells that respond, which remains low even at high doses. In contrast, the dTAG system enables the creation and manipulation of oscillatory protein expression dynamics, via targeted fine-tuned concentration-dependent degradation of our protein of interest, NGN3, homogenously across all cells in the population. By combining the dTAG and Tet-On systems, modifications of the existing TET-induced oscillations can be achieved, allowing for the creation of oscillations of periods shorter than the inherent TET-driven oscillations (shorter than 15 hours) in the cells that respond to doxycycline. However, expressing degron-tagged proteins via a constitutive promoter enables full temporal control of expression in all cells by controlling the timing of degradation and re-accumulation of the protein by the addition and removal of the drug, respectively. Pulsed application of dTAG drug allows us to create oscillations in protein expression over time, and we have named this system COD: **C**onstituitive promoter driving **O**scillations via **D**egradation. By utilising mathematical modelling we efficiently inferred the optimal timings for dTAG addition and removal to manipulate these COD-driven oscillations to create oscillations of different period lengths while maintaining the similar mean protein levels and peak-to-trough fold-changes. Therefore, we show that expressing degron-tagged proteins via a constitutive promoter permits precise, programmable control of protein expression dynamics.

## RESULTS

### Tet-On allows for dose dependent activation of oscillatory expression of unstable proteins of interest

To create oscillatory protein expression, we began by focusing on the first phase of the cycle: generating an expression peak. We turned to the Tet-On system and began by characterising the activation dynamics at the single-cell level over time in a clonal cell line. This system works via addition of doxycycline to the media, whereby the Tet-On transactivator binds the Tet3G promoter and expression of our protein of interest is initiated (Fig. 1A). We generated a clonal PANC-1 cell line containing the Tet3G system driving NGN3-mVenus-FKBP-V (NGN3-linker-3xFLAG-mVenus-linker-FKBP12^F36V^) expression, with a constitutively expressed NLS-mScarletI nuclear marker. We chose NGN3 as our protein of interest to test this system, since it has oscillatory expression endogenously (Miller et al. 2025) and there is a need to be able to manipulate the oscillations to understand their function. In addition, we used a direct fusion to mVenus fluorophore to be able to visualise NGN3 protein dynamics over time and the FKBP-V degron tag was added for later characterisation (see next section).

**Fig. 1.**
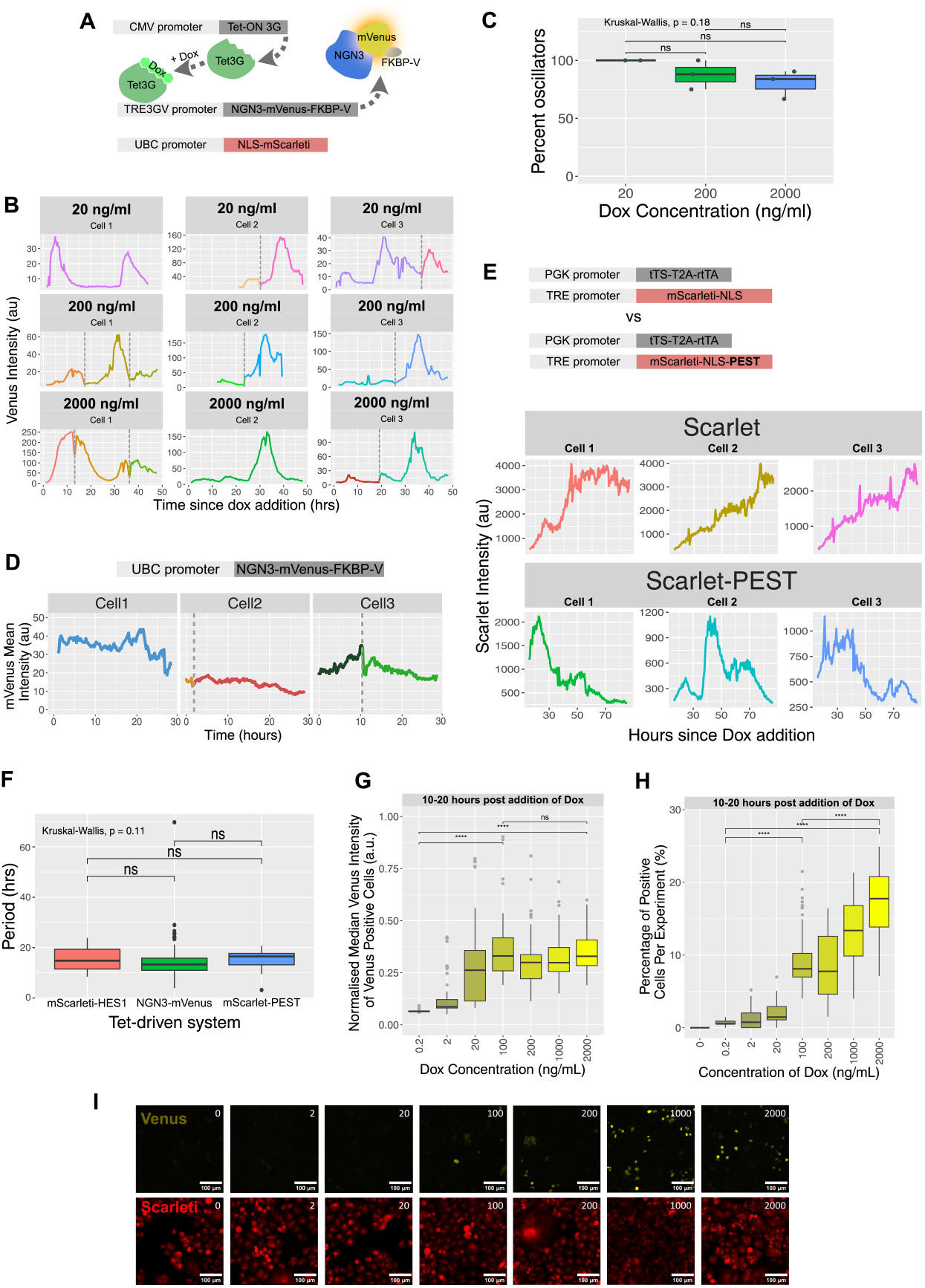
The Tet-On system allows for doxycycline induced activation of oscillatory expression of unstable proteins. (A) Schematic illustrating the constructs present in the clonal PANC-1 cell line (Tet-ON system driving NGN3-mVenus-FKBP-V plus NLS-mScarletI nuclear marker. Upon addition of doxcycline, the Tet-On 3G transactivator binds doxycycline, inducing transcriptional activation of NGN3-mVenus-FKBP-V via the TRE3GV promoter. (B) Single cell traces showing mean mVenus intensity over time in cells activated with 3 different concentrations of doxycycline. Grey dotted lines indicate cell division, with one daughter cell shown after division. Doxycycline concentration is indicated above each trace. (C) Percent of oscillatory mVenus positive cells at three different concentrations of doxycycline (dox). Points indicate independent experiments, n=3; Kruskal-Wallis p=0.18 (D) Single cell mVenus mean intensity over time in PANC-1 cells expressing NGN3-mVenus-FKBP-V from the constitutive UBC promoter. Grey dotted lines indicate cell division, with one daughter cell shown after division. (E) Single cell traces of MCF7 cells expressing either mScarletI::NLS or mScarletI::NLS::PEST driven by the Tet-On system. Cells were induced with 2000 ng/mL doxycycline. (F) Periodicity of cells that passed as oscillators expressing Tet-driven mScarletI::Hes1 (n=25), NGN3::Venus (n=162) or mScarletI::NLS::PEST (n=8) from 2-3 independent experiments. Kruskal-Wallis p = 0.11 (G) Median intensity of mVenus in positive cells (determined by mVenus expression higher than the maximum mVenus intensity of the 0ng/mL doxycycline control) 10-20 hours following addition of different concentrations of doxycycline (Dox). Median intensity levels per time point of mVenus are normalised to median intensity levels of the mScarletI nuclear marker. N=2-3 independent experiments, n=25-90 medians per boxplot. Kruskal-Wallis followed by Dunn posthoc test for multiple comparisons. (H) The percentage of mVenus positive cells (determined by mVenus expression higher than the maximum mVenus intensity of the 0ng/mL doxycycline control) 10-20 hours following addition of different concentrations of doxycycline (Dox). N=2-3 independent experiments, n=5971-22240 mVenus cells spread over 30 timepoints per boxplot. Kruskal-Wallis followed by Dunn posthoc test for multiple comparisons. (I) Images showing NGN3-mVenus expression (top row) and the same cells’ NLS-mScarletI expression (bottom row) at 20 hours post addition of doxycycline. Doxycycline concentration is displayed in the top right of each image in ng/mL. Scale bars = 100µm.

Unexpectedly, single-cell time-lapse imaging after doxycycline addition revealed Tet-ON driven NGN3::mVenus expression was highly dynamic (Fig. 1B). The majority of cells displayed multiple peaks and troughs over time, and more than 75% of cells passed our statistical oscillator test (Fig. 1C), showing oscillatory Venus expression regardless of doxycycline concentration. To validate that these dynamics were a result of the Tet-On system and not an inherent property of NGN3, we expressed NGN3 under a constitutive promoter (UBC) in PANC-1 cells (using a UBC-NGN3-linker-3xFLAG-mVenus-linker-ecDHFR-linker-FKBP12^F36V^ plasmid, where the addition of the destabilising domain, ecDHFR, was found to be non-functional as it had no effect on the protein stability (Fig. S1A) and is therefore not considered further). Under the UBC promoter, NGN3 expression was steady and sustained (Fig. 1D), and no cells pass our statistical oscillator test indicating that the oscillations were specific to the Tet-ON system.

To further test the Tet-driven spontaneous dynamics, we also examined a Tet-On system (2nd generation Tet system) in a different cell line (MCF7 cells), driving a different protein (mScarletI::HES1; Fig. S1B). Again, we observed dynamic protein expression following doxycycline addition (Fig. S1B), suggesting the dynamics could be caused by the Tet regulatory elements, and not an artifact arising from the PANC-1 cells. However, since both NGN3 and HES1 are also known to oscillate in their expression when expressed endogenously (Miller et al. 2025; Shimojo, Ohtsuka, and Kageyama 2008; Hirata et al. 2002; Sabherwal et al. 2021; Kobayashi and Kageyama 2010; Seymour et al. 2020; Weterings et al. 2024; Marinopoulou et al. 2021), we also generated additional MCF7 cell lines containing Tet-ON driven fluorescent proteins only (mScarletI). Dynamic expression was observed only when mScarletI was destabilised via a PEST domain, suggesting that protein half-life plays a key role, where long protein stability masks the dynamic properties of the Tet promoter (Fig. 1E). Interestingly, the oscillation periods of Tet-driven NGN3-mVenus-FKBP-V, mScarletI-HES1 and mScarletI-NLS-PEST were highly similar (a mean period of 13.9, 15.5 and 14.8 hrs respectively (Fig. 1F)), despite HES1 and NGN3 having distinct period lengths when expressed endogenously in human cells (Miller et al. 2025; Sabherwal et al. 2021; William et al. 2007; Cottrell et al. 2025). This suggests that the oscillatory protein expression observed is due to the Tet regulatory system rather than due to the protein expressed.

To further understand how the Tet-system creates oscillations in expression, we next asked if there was any link between the timing of the pulses and the cell cycle. We tracked single-cell expression dynamics of Tet-On driven NGN3-mVenus-FKBP-V or mScarletI-HES1 in PANC-1 or MCF7 cells respectively across mitoses. We found that Tet-On driven NGN3-mVenus-FKBP-V expression peaks could occur at any point during the cell cycle, while Tet-On driven mScarletI-HES1 peaks occured predominately at the start or end of the cell cycle (Fig. S1C,D). Therefore, this indicates that the Tet-On system is functional throughout the cell cycle, but it is advisable to characterise the Tet-driven expression dynamics for each protein of interest as the expression of the target protein can still be subject to cell-cycle dependent regulation, similar to HES1 (Hardwick and Philpott 2019; Cottrell et al. 2025).

Together, these findings indicate that the Tet-ON system can induce dynamic, oscillatory expression at single cell level, across multiple cell lines, Tet system generations, and target proteins (provided the expressed protein has a sufficiently short half-life).Thus, Tet-ON–driven expression may be harnessed to generate population-wide asynchronous protein oscillations at a fixed period, in contrast to constitutive promoters (e.g., UBC), which yield steady, sustained expression

In order to test the function of the different parameters of an oscillating signal (e.g., period, level, fold-change/amplitude, phase), it is essential to have precise control over each of these parameters. Therefore, we then asked whether Tet-driven oscillations could be engineered to allow systematic tuning of these parameters. Due to population studies that show an increase in protein expression with an increasing dose of doxycycline (for example Kato et al. 2025; Zhu et al. 2024; C.-H. Park et al. 2012), we hypothesised that changing the doxycycline concentration might enable us to have control over the level of the resulting protein oscillations. We added doxycycline at different concentrations (0.2, 2, 20, 100, 200, 1000, and 2000 ng/µL) before time-lapse live imaging and found that at single cell resolution the concentration of doxycycline had little effect in changing the expression levels: NGN3-mVenus-FKBP-V levels in 100 ng/µL doxycycline was not significantly different to levels in 2000ng/µL doxycycline from 0-30 hours post doxycycline addition (Fig. 1G, S1E). However, the number of mVenus positive cells increased with increasing doxycycline concentration at all time points, suggesting that transcriptional activation is dose dependent (Fig. 1H,I, S1F). The proportion of mVenus positive cells peaked between 10 and 20 hours post-addition of doxycycline at the highest concentration used with a maximum of around 25% of cells positive for mVenus (despite the fact that this is a clonal cell line where every cell contains the same transgene, Fig. S1F). Therefore, we find that at the single cell level, increasing the doxycycline concentration in the Tet-On system increases the number of cells that respond to doxycycline, but at concentrations above a threshold of 100 ng/µL, does not change the resulting level of expression of the gene activated.

Further characterisation of the Tet-driven NGN3-mVenus-FKBP-V expression dynamics at different concentrations of dox(20, 200, and 2000 ng/µL doxycycline) with statistical analysis of time-series data showed no difference in the quality of oscillations (using Log likelihood ratio (LLR); high scores represent a better fit to a periodic model and are therefore an indicator of a high-quality oscillator, Fig. S1G.). Likewise, there was no difference either in the period of oscillations between the conditions examined, with all concentrations having an average period of around 15 hours (Fig. S1H). While we did see a significant difference in the maximum fold change (measured as the ratio between peak and trough) between 200ng/µL and 2000ng/µL doxycycline, this result was inconsistent and not dose dependent (i.e. there was no different between 20ng/µL and 2000ng/µL doxycycline nor between 20ng/µL and 200ng/µL doxycycline, Fig. S1I). Overall, our findings suggest that although the Tet-ON system can induce gene expression and create the required expression peak, it is not experimentally tunable. Instead, we find that at the single cell level, the Tet-On system drives dynamic protein expression with a fixed period even with continuous dox administration and regardless of the doxycycline concentration or human cell-line used.

### The dTAG degron system allows for the generation of oscillatory protein expression through control of protein degradation as well as re-accumulation

Due to the limited tunability of Tet-ON–driven protein expression oscillations, we next examined whether incorporating the dTAG degron system could enable modulation of specific features of oscillations, or whether combining the degron system with a constitutive promoter would allow full control over protein dynamics. The dTAG approach relies on introducing a small epitope tag (FKBP12^F36V^) that, upon addition of a dTAG drug into the media, recruits endogenous degradation machinery to rapidly degrade the tagged protein (Fig. 2A). Several dTAG molecules have been developed, and in this study we focused on degradation using dTAG^V^-1, dTAG-13, and dTAG-47 after expression either by the dynamic Tet-ON promoter or constitutive UBC promoter.

**Fig. 2.**
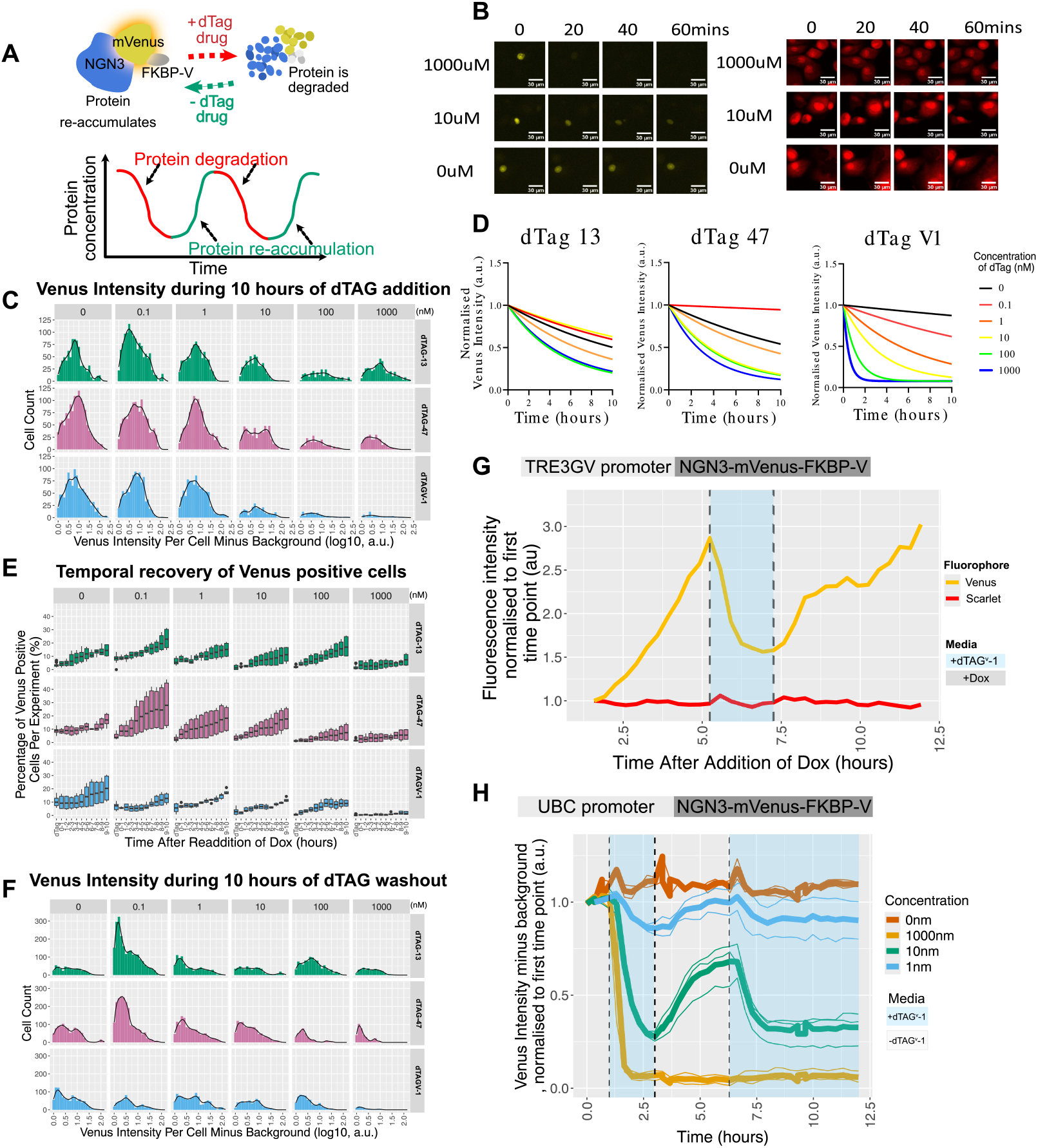
The dTAG degron system allows for the generation of oscillatory protein expression through control of protein degradation as well as re-accumulation. (A) the presence of the FKBP-V motif on a protein of interest allows dTAG drug to target the protein for degradation, and removing the dTAG drug from the system removes dTAG mediated protein degradation allowing re-accumulation of protein. The aim to be able to repeatedly degrade and re-accumulate in order to create oscillations of expression. (B) mVenus positive cells treated with dTAG^V^-1. Left images shows mVenus while right grid shows the corresponding NLS-mScarletI expression in the same cells. Time after addition of dTAG^V^-1 is displayed along the top of each grid while concentration of dTAG^V^-1 is displayed along the left. Scale bars = 30µM. (C) Density plot showing mVenus intensity across a population of cells treated with different concentrations of 3 different dTAG drugs for a duration of 10 hours (30 timepoints). mVenus intensities are background subtracted and only positive mVenus intensities were plotted. N=3 independent experiments, n=52-1319. (D) Regression plots of degradation of NGN3-mVenus-FKBP-V following addition of different dTAG drugs at different concentrations, using data from Fig. S2A. Data was derived from mean data of single cell mVenus intensity traces (n= 6-22) from two independent experiments. (E) The percentage of mVenus positive cells over a 10 hour time period after dTAG is removed and doxycycline is added. Data is faceted by concentration of dTAG (nM) on the x axis and the dTAG drug the cells were released from on the y axis. Data is binned by hour. The first bin (dTAG) is the last remaining 5 hours cells spent in dTAG drug treatment before media change out of drug. N=2 independent experiments, n = 9083 - 19893 cells used to generate percentages per panel. (F) Density plots showing the intensity of mVenus across the population of mVenus positive cells following release from 3 dTAG drugs at different concentration (x axis) over a period of 10 hours. N=2 independent experiments, n=78 (dTAG^V^-1, 1000nM) 2248 (dTAG-13, 0.1nM). (G) Mean intensity of mVenus (yellow) and mScarletI (red) for a population of cells normalised to time 0, sequentially exposed to 2000 ng/mL doxycycline, 10nM dTAG^V^-1 (blue shaded area), and re-addition of 2000 ng/mL doxycycline. Data is from a single experiment, n=25 cells. Single cell traces are in Fig. S2D. (H) Addition and removal of dTAG^V^-1 to PANC-1 cells expressing NGN3::mVenus::FKBP-V under a UBC promoter. Cells were periodically exposed to media containing different concentrations of dTAG^V^-1 (or DMSO as the control, labelled here as 0nm). The blue highlighted regions between the vertical dashed lines indicates the time in which cells were in dTAG^V^-1. Single cell traces are in Fig. S2E-H. Means are the mean intensity of a whole field of view at any given timepoint. N=4-5 experiments.

We aimed to create oscillations and therefore required two key features: controllable protein degradation (to establish expression troughs) and the ability to remove the drug to permit protein reaccumulation, with the cycle repeatable over multiple rounds to produce successive peaks and troughs (Fig. 2A). Therefore, we first examined the temporal dynamics of NGN3 degradation (after Tet-driven expression) using three different dTAG compounds in a range of concentrations, and secondly assessed protein recovery after drug washout.

Degradation was dose-dependent (Fig. 2B,C,D): higher concentrations of each compound drove rapid and near-complete loss of NGN3 expression, while lower concentrations reduced but did not fully eliminate fluorescence. Among the tested drugs, dTAG^V^-1 was the most effective, degrading NGN3 to undetectable levels both more rapidly and at lower concentrations than dTAG-13 or dTAG-47 (Fig. 2C,D, S2A). This effect was specific, as the nuclear marker NLS-mScarletI remained unaffected across all treatments (Fig. S2B).

We next asked how reversible degradation was across compounds and concentrations, testing whether NGN3 could reaccumulate after drug washout and doxycycline re-addition. The most effective degradation condition— 1000 nM dTAG^V^-1—was also the least reversible: even 10 hours after washout, mVenus expression was still largely undetectable (Fig. 2E), and the few cells that regained fluorescence exhibited very low intensity (Fig. 2F). By contrast, lower dTAG^V^-1 concentrations or the other dTAG compounds permitted more robust recovery, with a greater fraction of cells regaining NGN3 and at higher levels (Fig. 2E,F). As before, the nuclear marker (NLS-mScarletI) remained unaffected in all conditions (Fig. S2C). These results highlight a critical trade-off between efficient degradation and the ability to restore protein expression upon drug withdrawal.

With this in mind we created a dynamic expression pattern, creating a peak and trough in expression. We selected a maximal doxycycline concentration of 2,000 ng/µL to induce NGN3-mVenus-FKBP-V expression in as many cells as possible, and 10 nM dTAG1-^V^1 to degrade the protein - this concentration was effective at reducing protein levels and was also able to be washed out successfully (Fig. 2C,D,E,F). By pulsing cells first with doxycycline, then briefly with dTAG, and returning to doxycycline, we successfully produced a distinct peak, trough and subsequent peak in NGN3 expression, while the nuclear marker NLS-mScarletI remained unaffected by the treatments (Fig. 2G). These dynamics were clearly observable at the population level illustrating synchronicity in the population (Fig. 2G), but single-cell traces (Fig. S2D), however, revealed heterogeneous responses. For example some cells exhibited low expression until the second doxycycline pulse, while others peaked with the first but not the second pulse of doxycycline. Despite this variability, these results demonstrate that combining the Tet-ON and dTAG systems enables controlled synchronous oscillatory modulation of protein levels, allowing the creation of shorter periodicities than the inherent Tet-ON driven protein expression oscillations.

Finally, to assess whether promoter context affects responsiveness to dTAG pulses, and to overcome the variability seen in the Tet-driven promoter system, we tested constitutive UBC-driven NGN3-mVenus-FKBP-V. In contrast to the Tet system, all cells express NGN3 in the absence of dTAG, which creates a more homogeneous population of cells with sustained expression across all cells (Fig. 1D, 2H, S2E). Additionally, in the presence of dTAG all cells respond, and similar to the Tet-ON system, higher concentrations of dTAG1-^V^1 enhanced degradation efficiency but reduced reversibility (Fig. 2H, S2F,G,H). We generated a two-peak oscillatory expression pattern using this setup, and again identified 10 nM dTAG1-^V^1 as the optimal concentration, effectively degrading the protein while allowing robust recovery upon washout, thereby producing two clear pulses of expression (around 6 hours apart) when the dTAG drug was pulsed in and out of the media in all cells (Fig. 2H, S2F,G,H). In conclusion, combining a constitutive promoter with the dTAG system yielded a more uniform, homogenous response at the single cell level compared with the Tet-On system, allowing for the generation of dynamic protein expression in all cells. As such, we continued with this system, and named it COD: **C**onstituitive promoter driving **O**scillations via **D**egradation.

### The dTAG degron system enables control of oscillatory parameters from constitutively expressed proteins (COD system)

Our next goal was to manipulate protein expression dynamics to create oscillations with tunable parameters using the COD system (UBC-driven NGN3::mVenus-FKBP-V PANC-1 cell line), which exhibits more homogeneous expression across cells compared to the Tet-ON–driven system. Specifically, we sought to determine whether NGN3 protein oscillations could be produced with distinct periods, while keeping other oscillatory characteristics constant (levels and peak-to-trough fold-change). This would enable us to demonstrate the degree of control we can achieve over protein expression dynamics, and also isolate and investigate the specific role of NGN3’s period length, which is an aspect that remains poorly understood. To identify the optimal experimental setup to achieve this we turned to mathematical modelling. This approach was essential as balancing period changes with constant levels and fold-change is not intuitive, and it would require extensive trial and error, making the process both time-consuming and costly.

We constructed an ordinary differential equation model (see (1) in Methods) to quantitatively describe the temporal regulation of NGN3 under dTAG-induced degradation. The model assumes that NGN3 exists at a steady-state level in the absence of drug (as seen in Fig. 1D, 2H) due to constant and balanced basal production and degradation rates, and that NGN3 is then further degraded upon addition of the dTAG1-^V^1 drug. Using the data at 4 different dosages (0nm, 1nm, 10nm, 1000nm) from Fig. 2H we parameterised the model to fit kinetic parameters for basal production and degradation and the dependence of NGN3’s degradation rate on dTAG1-^V^1 drug (Fig. S3A). This allowed us to simulate different oscillatory scenarios and thus inform the optimal estimates for drug-scheduling times and dosage levels required for creating oscillations with differing periods. Accurate reproduction of the dynamics in Fig. 2H required modelling NGN3 degradation as a non-linear function of drug concentration (Fig. S3B) and accounting for gradual, rather than instantaneous, drug clearance (Fig. S3C).

As a proof of principle, we asked the model (with parameters now at fixed values) to infer the optimal drug-scheduling times required to generate NGN3 oscillations with 10-hour and 5-hour periodicities using 10nm of dTAG1-^V^1, while maintaining similar mean levels and fold-change/amplitude (Fig. 3A,B). To achieve this, model was fitted to test data: a synthetic, sinusoidal wave with tunable parameters (i.e. period, mean level, min/max values). We tuned these parameters to mimic waves with our desired oscillatory properties (i.e. 10-hour or 5-hour periods, yet sharing a fixed mean level and maximum and minimum values: see Test data in Fig. 3C-E and 3G-I for 5-hour and 10-hour sine waves respectively, with quantification of the Test data parameters in Fig. 3F,J) and then searched a range of drug-on/off time-window schedules, comparing the model simulation and test data at each. From this search, the optimal dTAG1-^V^1 addition and removal times were calculated to produce the best agreement between the model and the target oscillations (Fig. 3A,B). Intuitively, for the 10-hour periodicity, the model predicted that alternating 5 hours of dTAG treatment with 5 hours of washout provides one of the best-fitting regimes (Fig. 3A, D). However, to create the best fit for the 5-hour periodicity alternating 1.5 hours/3.5 hours (drug present/drug washout) was optimal (Fig. 3B,I). Other regimes created the required period but were not able to maintain similar mean levels or fold-changes in the oscillations, for example 2.5 hours of drug present with 2.5 hours of drug washout created the required 5 hour period, but the mean level dropped by 22% compared to the 1.5 hour present /3.5 hours washout schedule (Fig. 3C,E,F,G,H,J).

**Fig. 3.**
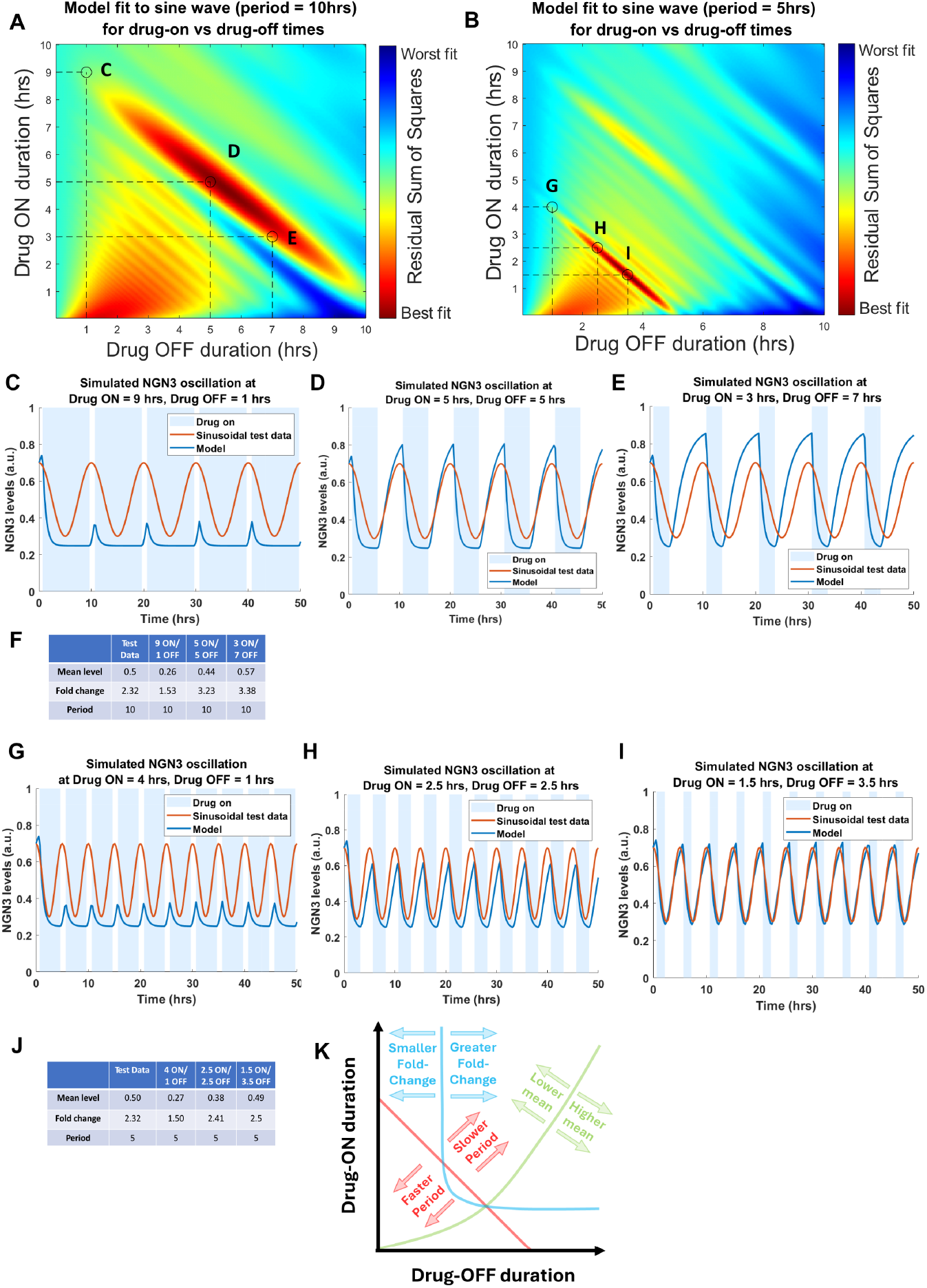
Inferring optimal drug pulsing timings to create desired oscillatory regimes. (A) Heatmap of sum of squares residual error between sinusoidal synthetic data with period of 10 hrs and model simulations with optimal parameters at drug concentration *d* = 10*nM* determined at varying repeating drug-ON/drug-OFF window times [0-10 hrs x 0-10 hrs]. (B) Heatmap of sum of squares residual error between sinusoidal synthetic data with period of 5 hrs and model simulations with optimal parameters at drug concentration *d* = 10*nM* determined at varying repeating drug-ON/drug-OFF window times [0-10 hrs x 0-10 hrs]. (C, D, E) Model solutions (blue lines) with drug concentration *d* = 10*nM*, taken at three different drug-ON/drug-OFF schedules (marked by black circles in panel A). Synthetic sinusoidal data (with period, mean level and peak-to-trough fold-change as shown in F) is indicated by the orange line. Blue shaded area indicates a drug-ON window. (F) Table showing properties (Mean level, peak-to-trough fold change and peak-to-peak period) of sinusoidal test data (orange waves in C, D, E) and model simulations (blue waves in C, D, E) at three different drug-ON/drug-OFF window times (marked by black circles in panel A). (G, H, I) Model solutions (blue lines) with drug concentration *d* = 10*nM*, taken at three different drug-ON/drug-OFF schedules (marked by black circles in panel B). Synthetic sinusoidal data with (with period, mean level and peak-to-trough fold-change as shown in J) is indicated by the orange line. Blue shaded area indicates a drug-ON window. (J) Table showing properties (Mean level, peak-to-trough fold change and peak-to-peak period) of sinusoidal test data (orange waves in G, H, I) and model simulations (blue waves in G, H, I) at three different drug-ON/drug-OFF window times (marked by black circles in panel B). (K) Schematic illustration of how adjustments in drug-ON vs drug-OFF durations perturb oscillatory properties of a wave (see Fig. S3D-F). Solid lines indicate points at which periodicity (red), fold-change (blue) and mean levels (green) remain invariant between a model (Eq. (1)) simulation at fixed parameters and a particular, arbitrary sinusoidal test vector. Corresponding coloured annotated arrows display the manner of perturbation of oscillatory property for a given ON/OFF timing adjustment in either direction away from each solid line.

To understand why the drug-on/drug-off schedules needed to achieve a target oscillatory profile do not necessarily align with intuitive expectations (e.g., symmetric on and off drug durations), we examined how varying these times affects the period, mean level, and peak-to-trough fold change of the oscillatory signal independently (Fig. S3D,E,F and also see schematic Fig. 3K). As expected, the period remains constant as long as the sum of the on and off durations matches the desired period length, becoming faster when both are short and slower when both are prolonged (Fig. 3K, S3F). The mean level shifts with the relative balance of on and off times: longer off phases compared to on generally yields higher mean levels, likely reflecting faster degradation upon drug addition than re-accumulation after drug removal (Fig. 2H, 3K, S3D). Finally, the model predicts that for the peak-to-trough fold-change, shorter on or off phases reduces the fold-change by limiting either degradation or recovery time. Interestingly, there is a point from which the system saturates, highlighting that if given enough time (to degrade, or re-accumulate) the expression of NGN3 will reach the steady state in the system (the expression level at which the protein is expressed in the absence or constant presence of the drug), and as such cannot change the fold-change further (Fig. S3E). In practice, this allows one to find optimal on/off timing at which the period, mean levels and fold change align with desired outcomes (illustrated by the three intersecting lines in Fig. 3K), which represent the areas of best fit on the heatmaps (Fig. 3A,B), also shown for periods from from 1hr to 16hrs (Fig. S3G-K).

To test the recommended drug-on/drug-off predictions from the model, we leveraged microfluidics, to alternate media containing 10 nM dTAG^V^1 with drug-free media at specific time intervals using our COD system (COD+CHIPS; Fig. 4A). By varying both the pulse duration and the interval between pulses, we successfully produced oscillatory waves of NGN3 at two distinct frequencies, compared to the control (sustained steady expression; Fig. 4B). Cells exposed to the shortest pulsing regime (1.5 hours drug on, 3.5 hours drug off) which was predicted to create a 5-hour periodicity in the model exhibited oscillations with an average 5.08 h period (Fig. 4B,C,D). Likewise, those under the longest regime (5 hours drug on, 5 hours drug off) predicted to create a 10 hour periodicity by the model, displayed a 9.33 hour period between the two peaks generated (Fig. 4B,C,D). In addition, as the model predicted, these pulsing regimes resulted in mean intensities and peak-to-trough fold-changes that were similar between the 5-hour and 10-hour periodicities created (Fig. 4C,D). Together, this shows that the model is able to accurately infer the optimal drug-scheduling times to generate oscillations with differing parameters, and that oscillatory protein dynamics can be flexibly tuned by adjusting dTAG pulse timing, providing a versatile framework for engineering synthetic oscillations.

**Fig. 4.**
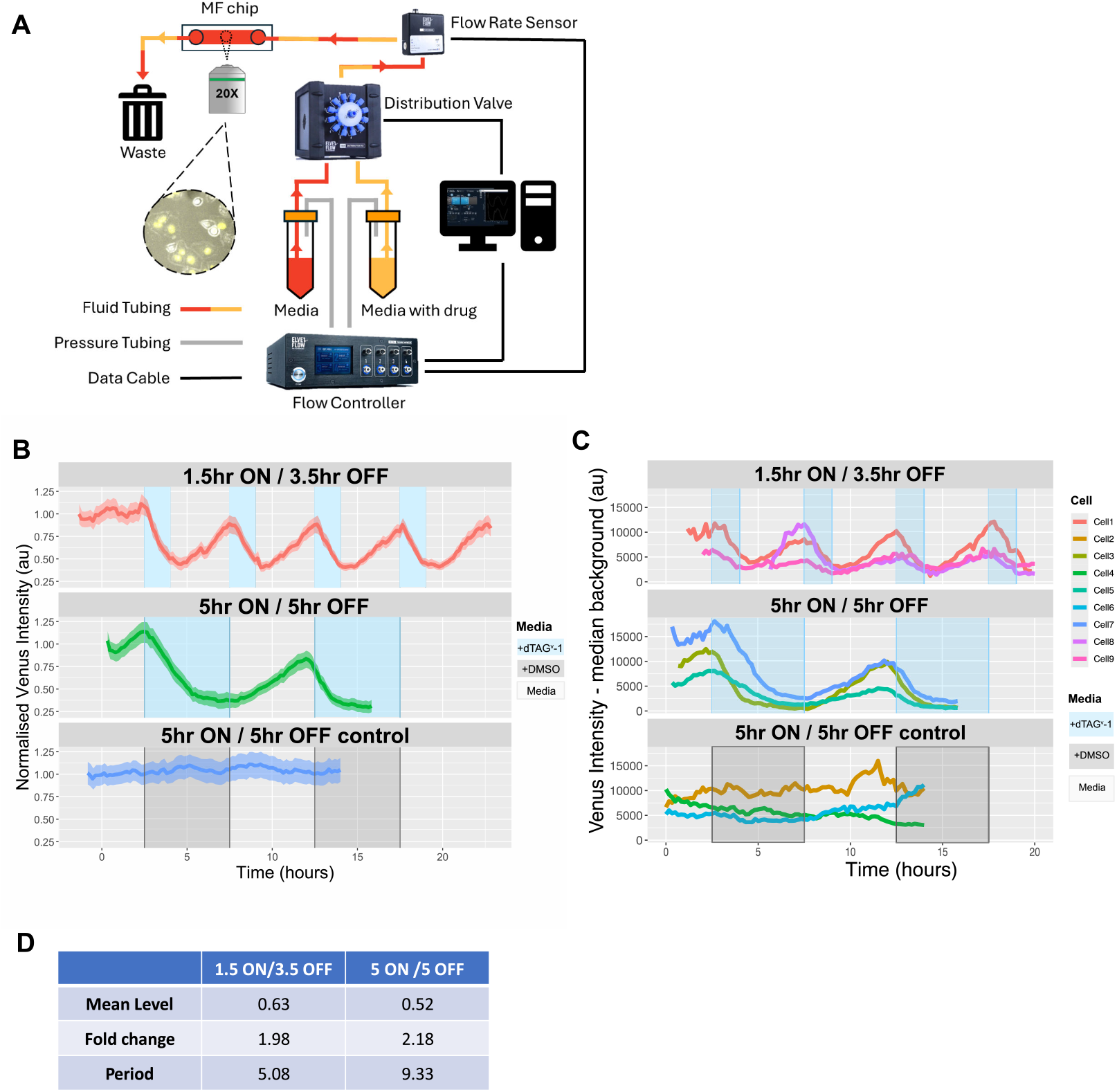
Creating desired dynamic regimes using the COD system and microfluidics (COD+CHIPS) (A) Diagram illustrating the microfluidic set up using Elveflow microfluidic instruments (more details in methods) to be able to switch between media +/−dTAG^V^-1 automatically. (B) Mean mVenus intensity levels of the entire field of view of UBC-NGN3-mVenus-FKBP-V cells, normalised to the intensities at the beginning of the time-lapse movie. Media containing 10 nM dTAG^V^-1 was applied to cells using microfluidics at different time intervals. To generate oscillations with a 5 hour period, cells were exposed to repeated intervals of dTAG^V^-1 for 1.5 hours (indicated by the blue highlighted areas) followed by 3.5 hours of media without the drug. To generate oscillations with a 10 hour period, cells were exposed to repeated periods of dTAG^V^-1 for 5 hours (indicated by the blue highlighted areas) followed by 5 hours without the drug. Control cells were exposed to media containing DMSO (indicated by grey highlighted areas) for 5 hours between periods of normal media. (C) Single cell examples of data from (B). Blue/grey highlighted areas indicate periods when cells were in dTAG^V^-1/DMSO. (D) Table showing properties (Mean level, peak-to-trough fold change and peak-to-peak period) of the experimental data in (B).

## DISCUSSION

Although both Tet-On and dTAG systems are widely used to induce gene expression or accurately degrade a protein of interest, respectively, there is insufficient characterisation related to reversibility and responsiveness at the single cell level. Therefore, here, using time-lapse single-cell microscopy, we show how these systems can be manipulated to achieve precise and temporal control of protein expression such that oscillatory expression of different characteristics can be created at will.

We find that the Tet-ON system produces protein oscillations in constant doxycycline administration, which is generated through a combination of transcriptional control (Tet-driven NGN3 has dynamic expression whilst UBC-NGN3 does not) and protein instability (Tet-driven mScarletI-PEST has dynamic expression whilst Tet-driven mScarletI does not). The existence of dynamics, particularly the troughs in expression whereby the protein is absent, explains other reports showing snapshot data of fewer cells expressing a Tet-on driven protein when the protein is destabilised (Pedone et al. 2019). We also observe stochasticity in the activation dynamics of the Tet promoter with only a minority of cells responding to doxycycline (25% of cells 10-20 hrs after dox), even in a clonal population when all cells are genetically identical. The stochasticity is also altered by doxycycline concentration, resulting in more cells responding at higher concentrations. Therefore, we believe population based measures showing increased expression in a dose-responsive manner is not due to each cell expressing more of the protein of interest, but rather more cells expressing the protein. Our data suggests that Tet-driven regulation happens in an all-or-none fashion, which is in agreement with other reports where saturation to doxycycline occurs at low concentrations (Pedone et al. 2019; Loew et al. 2010). This highlights the importance of studying single cell expression over time rather than snapshot-based population assays in order to draw biological conclusions from manipulations of any system.

Regardless of the unstable protein expressed via the Tet system, the periodicity was found to be approximately 15 hours under constant doxycycline administration. This indicates a potential refractory promoter state whereby the Tet promoter can not be activated in a time period shorter than 15 hours. Transcription is known to occur in bursts, switching between “on” and “off” states with the length of the “off” state highly influenced by the cisregulatory elements, chromatin architecture, transcription factor binding, PolII recruitment and the organism (transcriptional bursts in human cells occur on timescales of a few hours compared to a few minutes in flies; Suter et al. 2011; Cesbron et al. 2015; Lenstra et al. 2016; Lammers et al. 2020; Tantale et al. 2016). Since the Tet system comprises of two parts: the expression of the transactivator, followed by subsequent binding and activation to the TRE promoter in the presence of doxycycline, the 15 hour refractory period could be a combination of multiple refractory periods from both promoters, combined with sub-optimal activation. Indeed, optimising the promoter in which the transactivator is expressed depending on the cell lines used maybe be beneficial to create more homogeneity in the Tet-driven expression (Takiguchi et al. 2013).

We find that the dTAG degron system allows for a high degree of control over protein expression dynamics, especially when the protein is expressed constitutively (here using a UBC promoter, and named here as the COD system). A key advantage of this approach is its flexibility: any protein of interest can be directly targeted by fusing it with a degron tag, either endogenously (Abuhashem and Hadjantonakis 2022; Mehta et al. 2023) or exogenously, as in this study, where promoter choice can be tailored to the experimental context. It has also been shown that the addition of the dTAG epitope also does not affect the protein of interests function (Miller et al. 2025; Abuhashem, Lee, et al. 2022). Unlike expression-modulation systems that depend on manipulating upstream activators or repressors of signalling pathways (Sonnen, Lauschke, et al. 2018; Weterings et al. 2024), the dTAG system acts directly on the protein of interest. By contrast, signalling pathway-based modulation often involves affecting multiple steps in the signalling cascade before affecting the target protein, making it a relatively indirect and less specific strategy. Moreover, such approaches may inadvertently influence other downstream components of the signalling cascade, whereas the dTAG system avoids these off-target effects, and does not require prior knowledge of the regulatory mechanisms controlling a protein of interest. However, if upstream regulators or specific pharmacological inhibitors/activators are known then genetic engineering of the degron tag is not needed.

While other systems are also specific to the protein of interest, for example optogenetic regulation of expression (Isomura, Ogushi, et al. 2017; Shimojo, Isomura, et al. 2016; Imayoshi et al. 2013; Motta-Mena et al. 2014; Polesskaya et al. 2018) these rely on light activation which can have toxicity issues (Duke et al. 2020; Opländer et al. 2011; Wäldchen et al. 2015) and limits the fluorophores available for the experimental workflow (for example, a fluorophore that would be activated by the optogenetic light administration cannot be used as a downstream cell-fate reporter). In addition, while population-level light induction can be achieved, for example with led arrays, these are not often compatible with microscope setups (Kumar and Khammash 2022), therefore hindering simultaneous light activation and time-lapse fluorescent imaging. Instead, these optogenetic systems are ideal for localised induction, i.e. the activation of specific cells in a population. In contrast, the benefit of drug-based modulation approaches like dTAG, is the ability to synchronise the expression dynamics across a population of cells, while also enabling time-lapse imaging at single-cell resolution - achieving both high throughput and high resolution. This system can be further improved by the integration with microfluidics, in this study used to automate drug addition and removal, but in addition, can be used to study the dynamics of the micro-environment, for example mechanical forces and constraints, in a variety of embryos/tissue and organoids (S. E. Park, Georgescu, and Huh 2019; Sonnen and Merten 2019).

To finely-tune expression and create oscillations of different period lengths we varied the time interval of dTAG^V^-1 drug administration. Additional control can be achieved by modulating drug concentration and by selecting different dTAG drugs: both dTAG13 and dTAG47 recruit the CRBN E3 ligase complex while dTAG^V^-1 recruits the von Hippel-Lindau (VHL) E3 ligase complex (Nabet, Ferguson, et al. 2020; Nabet, Roberts, et al. 2018). This flexibility allows researchers to tailor drug choice to specific experimental contexts and protein targets, thereby enabling precise tuning of expression dynamics across diverse systems. Furthermore, the dTAG system can be integrated with complementary technologies such as optogenetics to achieve even finer temporal or spatial control. For instance, optogenetic gene activation precisely regulates transcription but does not influence protein degradation; thus, for highly stable proteins, achieving troughs in expression would require an additional mechanism such as dTAG-mediated degradation. Together, these features make the dTAG system a versatile platform for precisely controlling protein dynamics and dissecting the function of protein expression dynamics.

Through a mathematical modelling framework, we found that predicting how to achieve desired dynamic protein behaviours through drug pulsing is far from obvious. We included known experimental data for our protein of interest (protein half-life) and then identified model parameters that faithfully recapitulate experimental dynamics. Holding these parameters fixed, we discovered that the drug-on/drug-off schedules required to generate a target oscillatory profile do not necessarily align with intuitive expectations (e.g. symmetric on and off durations, each comprising half of the desired period). While a simple approach of repeating a pulse schedule whose drug-on+drug-off cycle matches the desired period can indeed produce oscillations of that period, this strategy can substantially perturb other key oscillatory features, such as peak-to-trough fold-change and mean expression levels. Instead, we found through computational exploration that optimal drug-timing regimes exist that must be followed to simultaneously control period, peak-to-trough fold-change, and mean levels. These findings underscore the value of computation in our approach; modelling here provides predictive power that refined experimental designs and revealed drug-timing schedules that would be difficult and costly to identify empirically, and provides a framework which can be used for any protein of interest.

## MATERIALS AND METHODS

For Materials and Reagents, see Table 1.

**Table 1.**
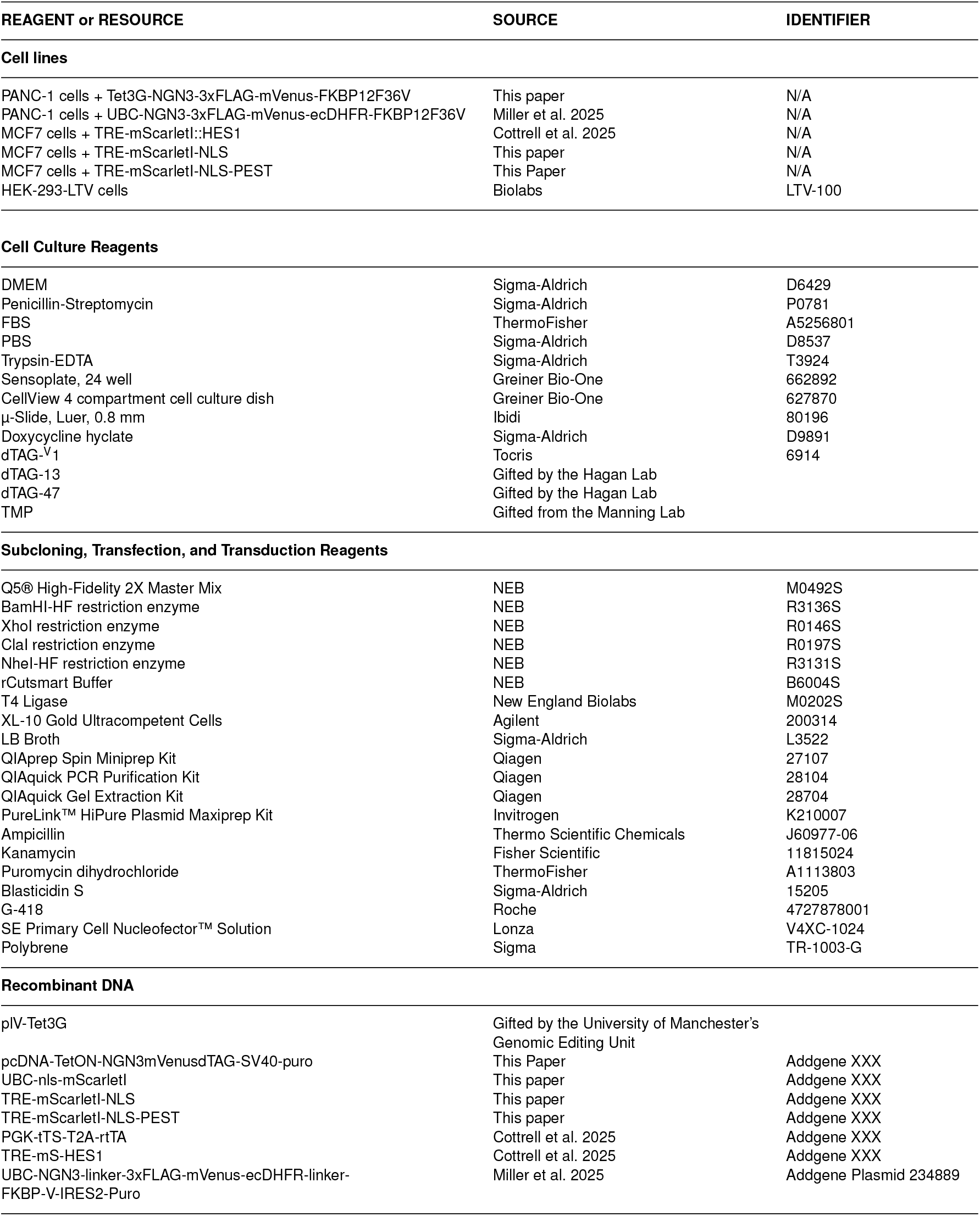
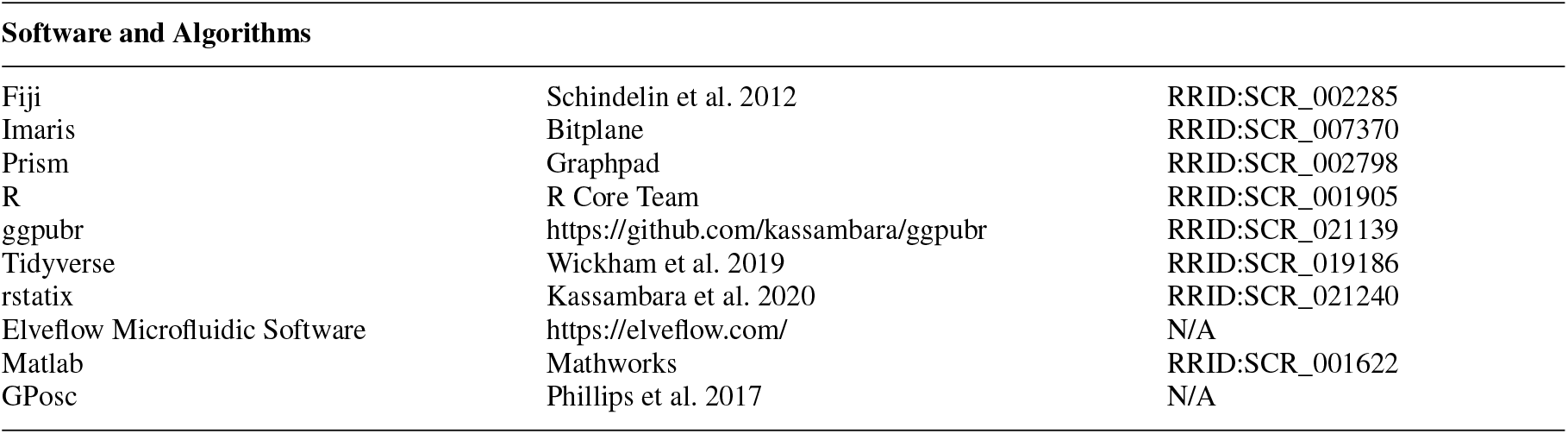
Materials and Reagents.

### Cell culture and cell line generation

PANC-1 and MCF-7 cells were cultured in high glucose DMEM (+ 10% FBS + 1x Pen/Strep) and maintained at 5% CO_2_ at 37°C in a humidified incubator. Routine Mycoplasma testing was carried out. PANC-1 cells were kindly gifted to us by Prof. K. Piper-Hanley. MCF7 cells were originally purchased from ATCC (HTB-22).

To generate our TetON-NGN3-mVenus-FKBP-V cell line, 1×10^6^ parental PANC-1 cells were nucleofected first with 2µg of the pcDNA-TetON-NGN3mVenusdTAG-SV40-puro plasmid (generated by the University of Manchester’s Genome Editing Unit (GEU)) and selected for with puromycin (2µg/mL). 1×10^6^ of the resulting cells were nucleofected with 2µg of the pLV-Tet3G plasmid (provided by the University of Manchester’s GEU) and selected for with blasticidin (3µg/mL). Nucleofections were performed using the Lonza 4D-Nucleofector®X Unit alongside the Lonza SE Cell Line 4D-Nucleofector®X Kit, pulse code DN100. Double nucleofected cells were transducted with lentivirus containing the UBC-NLS-mScarletI nuclear marker. The UBC-NLS-mScarletI plasmid was constructed by amplifying mScarletI using primers 5’-gcaggctagcg-ccaccatgccaaaaaagaagagaaaggtagtgagcaagggcgaggcag-3’ (fw) and 5’-gcagggatccttacttgtacagctcgtccatgccg-3’ (rev). The resulting mScarletI was ligated into pLNT-UBC-NLS-mVenus (Sabherwal et al. 2021) using T4 liagase following a restriction digest to remove the mVenus. The newly constructed plasmid was transformed into XL10-Gold ultracompetent cells (agilent) and a PCR colony screen was carried out. Promising clones were miniprepped and a restriction digest was performed as a further screen before sequencing. Following maxiprep the plasmid was packaged into lentivirus using HEK-293-LTV cells as described previously (Cottrell et al. 2025). PANC-1 cells were seeded at 1×10^5^ cells per well in a 6 well plate the day before exposure to virus. Cell media was changed with 1 mL of media containing 0.1% polybrene and virus was added and incubated with cells for 24 hours. Clonal lines were generated using FACS, with the activation of NGN3::mVenus expression by 2µg/mL doxycycline, and single cells positive for both mScarletI and mVenus fluorescence were sorted into 96 well plates. One clonal line was used for this study.

The Tet-ON driven mScarletI-NLS and Tet-ON driven mScarletI-NLS-PEST plasmids were constructed by amplification by PCR of the mScarletI gene with primers 5’-ggacggatc-ctctagagccaccatggatatcgtgagc-3’ (fw) and 5’-gcacctcgagttaatcga-ttacctttctcttcttttttgggtccttgtacagctcgtccatgcc-3’ (rev), and the PEST domain sequence using primers 5’-gcaggaggagcatgcatggtagcc-aatatcgatagccatggcttcccgc-3’ (fw) and 5’-tgcgattacgacgttgac-catgctagactcgagttacacattgatcctagcagaagcacag-3’ (rev). The TRE-NLS-mScarletI plasmid vector was digested with restriction endonucleases (NEB) to remove NLS-mScarletI to be replaced with mScarletI-NLS and mScarletI-NLS-PEST via T4 liagase (NEB) mediated ligation. The newly constructed plasmids were processed and screened as described above. Following maxiprep, the plasmids were packaged into lentivirus using HEK-293-LTV cells and transducted into previously generated MCF7 cells stably expressing the Tet-On transactivator as described above and in Cottrell et al. 2025. Cells positive for viral integration were selected for using 5 mg/mL G-418. Clonal lines were not generated for this cell line.

The MCF7-TetOn-Hes1-mScarletI cells were generated previously (Cottrell et al. 2025). PANC-1 cells expressing UBC-NGN3-mVenus-DHFR-FKBP-V were also generated previously (Miller et al. 2025). This cell line was previously sorted into 3 new polyclonal lines based on fluorescence levels of mVenus (high, medium and low expression, as seen in Miller et al. 2025). For this study, the high mVenus expressing cells were used. To test the function of the ecDHFR domain, TMP at 10µM was added to the cells and the Venus intensity was measured after 17 hours to assess protein levels.

### Timelapse imaging and Microfluidics

For experiments requiring more than four conditions, 1.2×10^5^ cells per well were seeded the day before imaging into Sensoplate black 24 well plates (Greiner-bio). Multiposition timelapses were generated using the Zeiss Celldiscoverer-7 and GaAsP-Pmt detectors by taking an image at each position every 20 minutes.A 20x 0.95 Plan-Aprochromat objective was used in combination with the Optovar 0.5x Tubelens. The mScarletI fluorophore was excited with a wavelength of 561nm and an emmision wavelength of 450-700nm was collected. The mVenus fluorophore was excited with 488nm and an emmission wavelength of 400-565nm was collected. Total resolution was 1661×1665, 1 pixel = 0.23µm, pinhole was approximately 2AU. Autofocus was utilised and a single Z per position was acquired per position for analysis. For experiments with 4 or fewer conditions, cells were seeded the day before imaging at 0.75-1×10^5^ cells into each quarter of a cellview 35mm 4 quarter dish (Greiner-Bio). Timelapses were generated using the Zeiss LSM880 microscope, GaAsP detectors and a Plan-Apochromat ×20 0.8 NA objective. Z-sections covering the depth of the cells were acquired every 10-20 minutes depending on experiment. Any manual media changes were performed using media equilibrated in 5% CO_2_ at 37°C for at least 1 hour. For experiments using microfluidics, the Elvesys Elveflow microfluidics system was used. Two media reservoirs were connected to a MUX microfluidic distribution valve and pressurised using the OB1 flow controller. The flow rate was detected by the MFS flow sensor. which feeds back to the OB1 (See Fig. 4A). Cells were seeded at 2.5×10^5^ cells/mL the day before imaging into a 0.8mm height single channel luer-lock µ-slides (Ibidi) according to the manufacturers protocol. The cells were imaged under flow using the Zeiss LSM880 microscope and timelapse videos were generated. Flow settings were set to 15µL-20µL/min throughout and the system was programmed using the Elveflow Microfluidic software.

### Cell Tracking

Z-stack movies were converted to max-projections based on the brightest pixel function in Fiji (RRID:SCR_002285). Whole-cell population data was generated using the ‘Spots’ function in Imaris (RRID:SCR_007370). Automatic nuclear detection using spots of 12µm in diameter was subsequently manually curated to ensure correct tracking. The ‘Spots’ function was also used to manually create single cell tracks with the ‘Track over time’ function in order to track and link values from the same nuclei over multiple frames. Mean intensity of mScarletI and mVenus for each spot was exported for further analysis.

### Oscillation analysis

To detect oscillations and to estimate period duration (amount of time it takes for one complete oscillation, e.g., from peak to peak) in our single-cell traces we used a customisable Gaussian pipeline described previously in Phillips et al. 2017 & Manning et al. 2019). The specific pipeline we used, as well as subsequent oscillation analysis, was described previously in Miller et al. 2025. The log likelihood ratio for each mVenus fluorescence single cell trace was determined by detrending the data and comparing it to Gaussian processes with both a wave-like covariance and a covariance with aperiodic fluctuations. The relationship between the probability of both defines the LLR which in turn determines whether a trace is considered oscillatory or non-oscillatory, following a false discovery rate test.

To calculate peak to trough fold-change we first defined the peaks and troughs in our mVenus fluorescence single cell traces using the Hilbert transform to generate instantaneous phase angles. We found the intensity values where each phase angle intercepted 0, giving us our peak and trough values which we then used to generate the peak to trough fold-change ratio. To track mVenus fluorescence across mitoses we used tRecs (https://github.com/TMinchington/tRecs), a custom python code which uses location, time, and fluorescence data to link traces either side of mitotic events, allowing for multi-generational cell tracking.

### Statistical Analysis

Statistical testing was performed in either GraphPad Prism 10.3.0 or R v.4.4.3 using the ggpubr or rstatix packages. Data was tested for normality using Shapiro-Wilk to select the appropriate test and significance was defined as ns p>0.05, *p<0.05, **p<0.01, ***p<0.001, ****p<0.0001. Statistical testing information for main figures including description, dispersion type, sample size and significance levels are included in figure legends. Statistical description of supplementary data is provided in supplementary figure legends.

### Mathematical modelling

#### Model description

NGN3 protein dynamics were described by the following first-order ordinary differential equation:

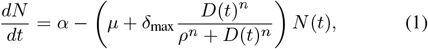

where *α* and *µ* denote the basal production and degradation rates of NGN3, respectively. In the absence of drug, the model assumes that NGN3 levels (represented by *N* (*t*) at time *t*) are governed by first-order production and degradation kinetics and predicts a steady-state protein level *N* ^∗^ = *α/µ*. Drug-induced degradation is represented by a Hill-type term that saturates with increasing drug concentration (illustrated in Fig. S3B). Here, δ_max_ is the maximal drug-induced degradation rate, *ρ* is the half-maximal effective drug concentration, and *n* is the Hill coefficient. *D*(*t*) denotes the time-dependent drug concentration. To reproduce experimental pulsing protocols, the drug exposure profile: Eq. (2), was modelled as a sequence of ON/OFF intervals determined by two parameters: the drug-ON duration (*T* ^*OFF*^ *i* − *T*^*ON*^ *i*) and drug-OFF duration (*T*^*ON*^ (*i* + 1) − *T*^*OFF*^ *i*), where *T*^*ON*^ *i* and *T*^*OFF*^ *i* denote the time points (hours) when the drug is applied and removed respectively, and *i* ∈ ℕ. During each ON interval, the drug concentration instantaneously rises to and remains at a fixed dose *d* (nm), and upon withdrawal, is removed according to first-order exponential decay:

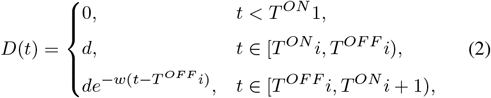

where *w* is the drug washout rate. Parameters *α, µ*, δ_max_, *ρ, n, d*, and *w* are assumed to be constants. A schematic illustrating how the drug levels *D*(*t*) behave temporally over a series of drug-ON/drug-OFF duration windows is provided in Fig. S3A.

#### Model Parameterisation

Model parameters were estimated by fitting solutions to Eq. (1) to mean experimental NGN3 time-course data collected under varying drug concentrations (0 nM, 1 nM, 10 nM, and 1000 nM) as seen in Fig 2 H. Experimental fluorescence values were first normalised by the maximum expression level across all doses and the model was initialised at the normalised NGN3 level at *t* = 0 for each dose. The drug exposure windows were set to match the experimental window times. Parameter fitting was performed using the nonlinear least-squares optimiser *lsqnonlin* in MATLAB where the objective function minimised the sum of squared residuals between model simulations (solved numerically using *ODE45*) and experimental trajectories across all four drug doses simultaneously with parameter *d* = [0, 1, 10, 1000], as seen in Fig. S3A. Initial estimates for parameters *α, µ*, δ_max_, *ρ, n*, and *w* were derived from biological expectations with realistically plausible bounds: degradation rate parameter *µ* was initialized 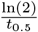 where *t*_0.5_ is the estimated NGN3 half-life which we took as 30 mins (based on Miller et al. 2025). Production rate *α* was then initially estimated as the ratio between the perceived steady state in the 0*nm* (no-drug) case and *µ*. The initial value for δ_max_ can be estimated as Deg_max_ − *µ*, where Deg_max_ is the largest rate of decrease under drug at the highest dosage, measured from the data (yellow line in Fig 2H). The half-maximal effective drug concentration *ρ*, was initially set as 5nm but could plausibly be estimated to be up to 20nm from the degradation data observed in Figs 2D and 2H. With little further real-world information to go on, the parameters *n* and *w* were estimated as integers of low-value. Conducting this process with time-course data from our first repeat resulted in the following set of parameters: {*α, µ*, δ_max_, *ρ, n, w*} ={0.42, 0.46, 3.30, 15.12, 1.37, 2.00}, which was then fixed as we proceeded to the drug exposure window exploration. Repeating this process across full experimental repeats yielded similar parameter values (the set of average parameter values across repeats were: {*α, µ*, δ_max_, *ρ, n, w*} = {0.425, 0.48, 2.65, 16.41, 1.285, 1.00}).

#### Drug window time exploration

To identify the optimal drug-exposure window regime required to yield oscillatory NGN3 at a desired period, we simulated model solutions at all repeating drug-ON and drug-OFF time window possibilities within a gridded range of 0-10 hours (with a mesh size of 0.1 hours). These model solutions were simulated with the fixed parameters found in the earlier model parameterisation, with *d* = 10, and compared (via a residual sum of squares (RSS) calculation) to synthetic sinusoidal Test data at a chosen period. Initial condition *N* (0) for the model was set to the value of the synthetic data *t* = 0. The optimal drug-ON time vs drug-OFF time for a wave of that chosen period was then determined by selecting the point at which comparison yielded the smallest RSS value.

### Use of Artificial Intelligence tools

During the preparation of this work, the authors used ChatGPT to reduce word count. After using this tool, the authors reviewed and edited the content as needed and take full responsibility for the content of the published article.

## Acknowledgements

The authors would like to thank the Bioimaging facility (especially Dr. James Bagnall and Dr. David Spiller), the Genome Editing Unit (especially Dr. Hayley Bennet and Dr. Antony Adamson) and the Flow Cytometry facility (especially Dr. Gareth Howell) of the University of Manchester. In addition, we would like to thank Prof. K. Piper-Hanley, Dr. Eleanor Trotter, Prof Iain Hagan, and Dr. Cerys Manning for aiding with provision of cell lines and reagents. We would also like to thank Dr. Cerys Manning and Dr. Antony Adamson for reading and providing comments on the manuscript, and all Papalopulu lab members for comments and discussions.

## Contribution

Conceptualization: N.P., A.M., Methodology: A.M., Software: A.R., Formal analysis: B.N., A.R., V.B., A.M., Investigation: B.N., O.C., A.R., F.W., X.W., A.M., Resources: A.M., Writing-original draft preparation: B.N., A.R., N.P., A.M., Visualisation: B.N., A.R., A.M., Supervision: N.P. and A.M., Funding acquisition: N.P. and A.M.

## Data availability

Code generated in this work can be found on https://github.com/AndyRowntree/Code-for-Noble-et-al, plasmids generated in this study have been deposited at Addgene XXX, and imaging data can be found at Figshare XXX

