## Supplementary info for "Engineering protein expression dynamics with Tet-ON and dTAG degron systems: from precise control to oscillations"

Fig.S1

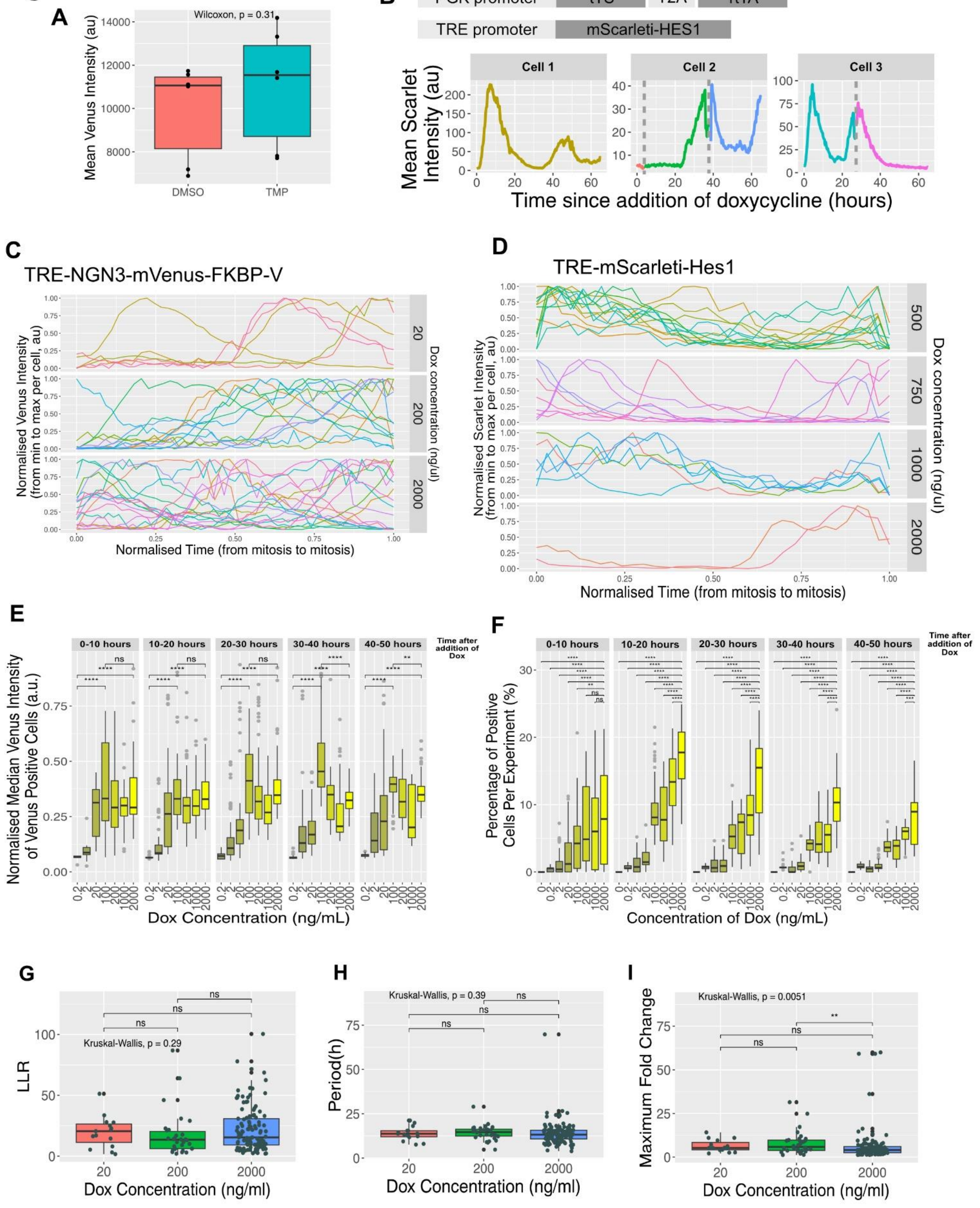
A
Wilcoxon, p = 0.31
Mean Venus Intensity (au)
DMSO
TMP
B
PGK promoter
tTS
T2A
rtTA
TRE promoter
mScarleti-HES1
Cell 1
Cell 2
Cell 3
Mean Scarlet Intensity (au)
Time since addition of doxycycline (hours)
C
TRE-NGN3-mVenus-FKBP-V
Normalised Venus Intensity (from min to max per cell, au)
Dox concentration (ng/ul)
Normalised Time (from mitosis to mitosis)
D
TRE-mScarleti-Hes1
Normalised Scarlet Intensity (from min to max per cell, au)
Dox concentration (ng/ul)
Normalised Time (from mitosis to mitosis)
E
0-10 hours
10-20 hours
20-30 hours
30-40 hours
40-50 hours
Time after addition of Dox
Normalised Median Venus Intensity of Venus Positive Cells (a.u.)
Dox Concentration (ng/mL)
F
Percentage of Positive Cells Per Experiment (%)
Concentration of Dox (ng/mL)
G
Kruskal-Wallis, p = 0.29
LLR
Dox Concentration (ng/ml)
H
Kruskal-Wallis, p = 0.39
Period(h)
Dox Concentration (ng/ml)
I
Kruskal-Wallis, p = 0.0051
Maximum Fold Change
Dox Concentration (ng/ml)

**Fig.S2**

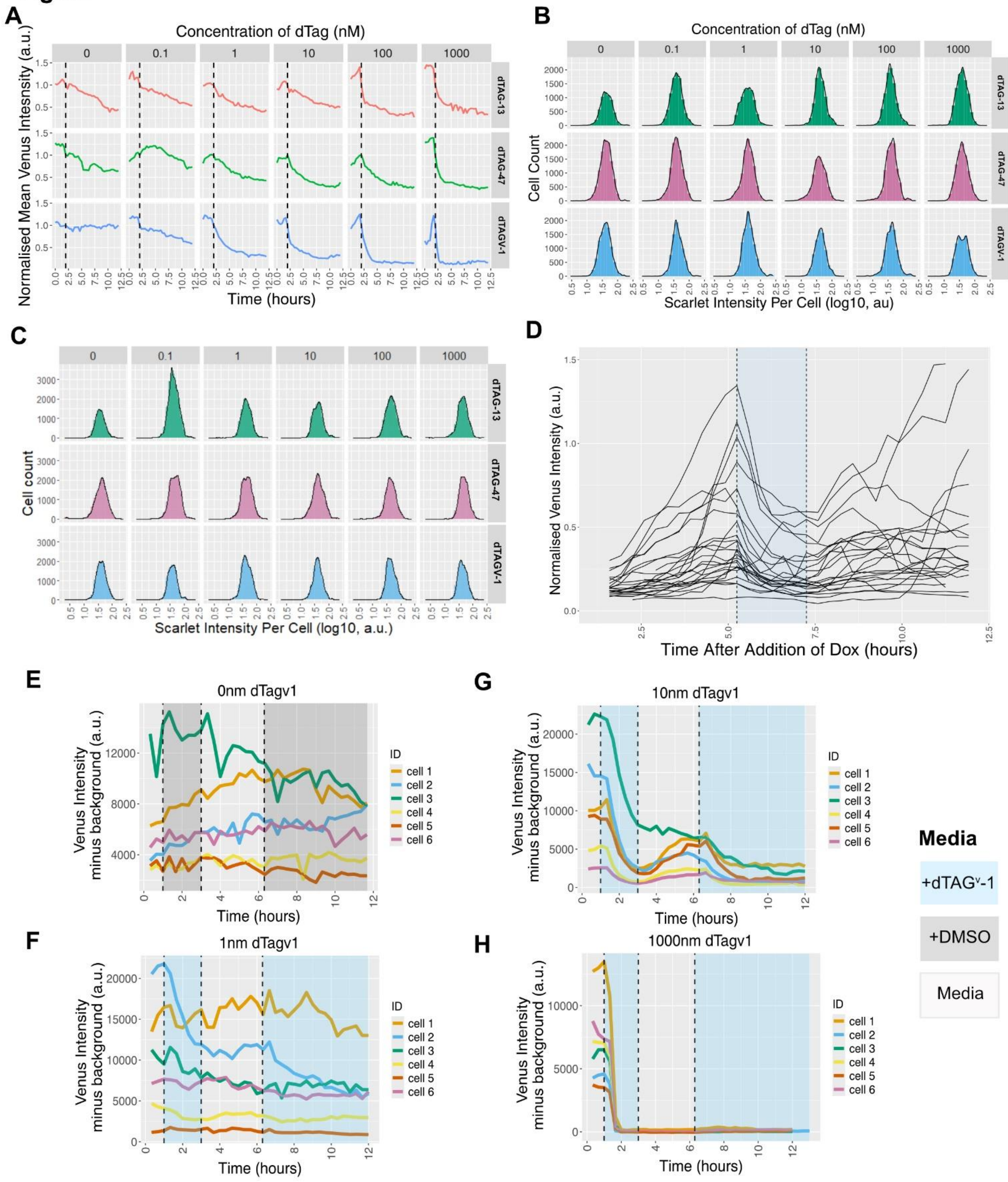
A
Concentration of dTag (nM)
0
0.1
1
10
100
1000
dTAG-13
dTAG-47
dTAGV-1
Normalised Mean Venus Intesnsity (a.u.)
Time (hours)
B
Concentration of dTag (nM)
Cell Count
Scarlet Intensity Per Cell (log10, au)
C
Cell count
Scarlet Intensity Per Cell (log10, a.u.)
D
Normalised Venus Intensity (a.u.)
Time After Addition of Dox (hours)
E
0nm dTagv1
Venus Intensity
minus background (a.u.)
Time (hours)
ID
cell 1
cell 2
cell 3
cell 4
cell 5
cell 6
F
1nm dTagv1
G
10nm dTagv1
H
1000nm dTagv1
Media
+dTAGᵛ-1
+DMSO
Media

### Fig.S3

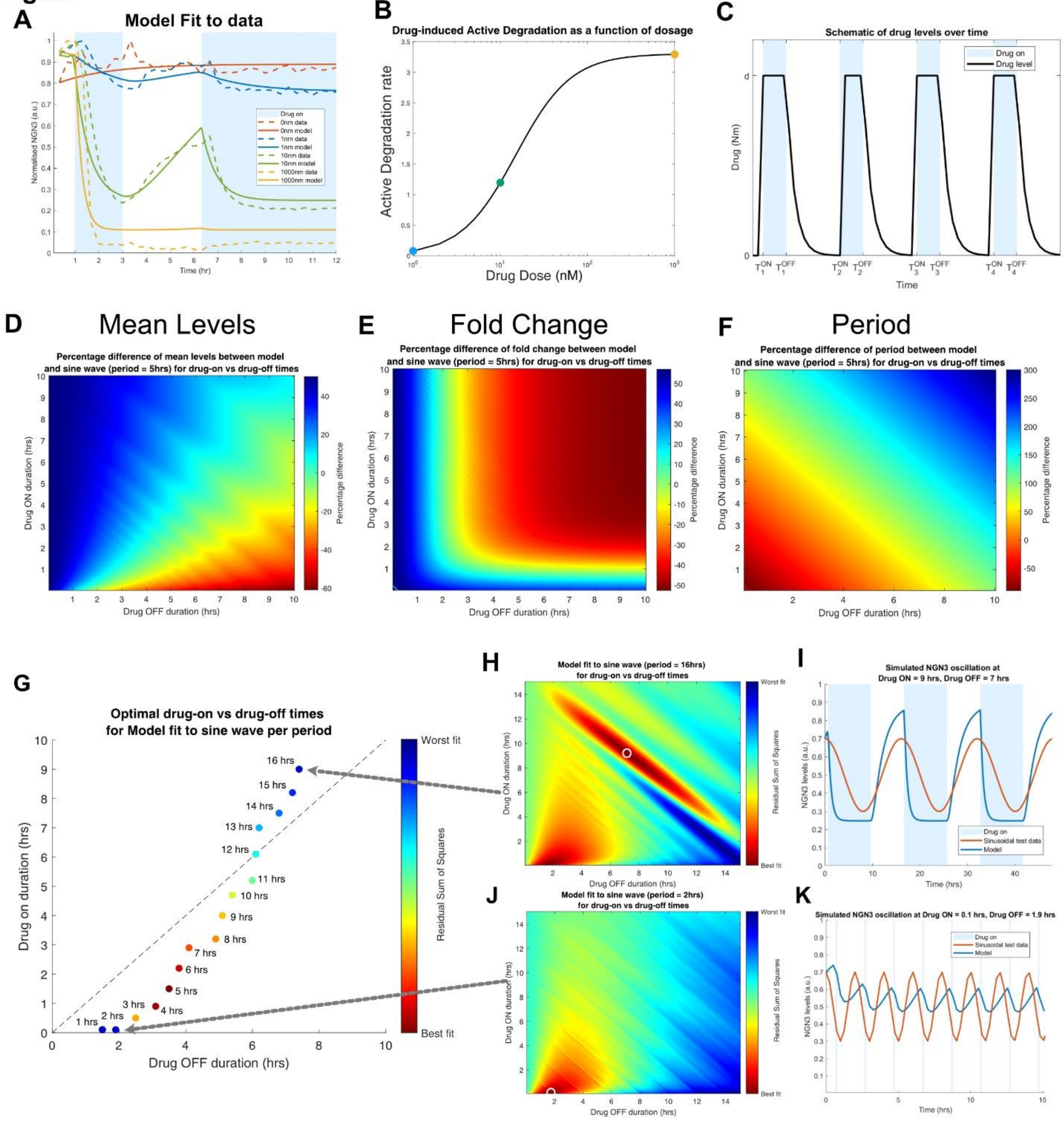

A
Model Fit to data
Time (hr)
Normalised NGN3 (a.u.)
Drug on
0nm data
0nm model
1nm data
1nm model
10nm data
10nm model
1000nm data
1000nm model
B
Drug-induced Active Degradation as a function of dosage
Active Degradation rate
Drug Dose (nM)
C
Schematic of drug levels over time
Drug on
Drug level
Drug (Nm)
Time
D
Mean Levels
Percentage difference of mean levels between model and sine wave (period = 5hrs) for drug-on vs drug-off times
Drug ON duration (hrs)
Drug OFF duration (hrs)
Percentage difference
E
Fold Change
Percentage difference of fold change between model and sine wave (period = 5hrs) for drug-on vs drug-off times
F
Period
Percentage difference of period between model and sine wave (period = 5hrs) for drug-on vs drug-off times
G
Optimal drug-on vs drug-off times for Model fit to sine wave per period
Drug on duration (hrs)
Drug OFF duration (hrs)
Worst fit
Residual Sum of Squares
Best fit
1 hrs
2 hrs
3 hrs
4 hrs
5 hrs
6 hrs
7 hrs
8 hrs
9 hrs
10 hrs
11 hrs
12 hrs
13 hrs
14 hrs
15 hrs
16 hrs
H
Model fit to sine wave (period = 16hrs) for drug-on vs drug-off times
I
Simulated NGN3 oscillation at Drug ON = 9 hrs, Drug OFF = 7 hrs
NGN3 levels (a.u.)
Time (hrs)
Sinusoidal test data
Model
J
Model fit to sine wave (period = 2hrs) for drug-on vs drug-off times
K
Simulated NGN3 oscillation at Drug ON = 0.1 hrs, Drug OFF = 1.9 hrs

**Supplementary Figure Legends**

**Figure S1. Characterisation of the Tet-ON expression dynamics at single cell level over time. Related to Figure 1.**

(A) Mean intensity of cells expressing UBC driven NGN3-mVenus-ecDHFR-FKBP-V expression after addition of 10µM TMP or DMSO for 17hours. n= 101-136 cells from 2 experiments. Wilcox test p= 0.31.

(B) Schematic of the second-generation TET-ON system tested in MCF7 cells driving mScarleti-HES1 and single cell examples of TET-ON driven mScarleti-HES1 expression after activation with 50ng/µl doxycycline. Grey dashed lines represent mitosis where afterwards one daughter cell is displayed.

(C) Single cell analysis of when the peaks in TET-ON driven NGN3-mVenus-FKBP-V expression occurs during the cell cycle for different concentrations of doxycycline (each panel). Cells were chosen which have similar lengths in their cell cycle (median cell cycle length +/- 0.5x standard deviation of the total population), and the normalised time represents time from mitosis to mitosis (0 and 1 represent the time of mitoses). The mVenus intensity is min-max normalised for each cell, and each line represents a single cell trace over time (n=6-18 cells from 3 independent experiments).

(D) Single cell analysis of when the peaks in TET-ON driven mScarleti-HES1 expression occurs during the cell cycle for different concentrations of doxycycline (each panel). Cells were chosen which have similar lengths in their cell cycle (median cell cycle length +/- 0.5x standard deviation of the total population), and the normalised time represents time from mitosis to mitosis (0 and 1 represent the time of mitoses). The mScarleti intensity is min-max normalised for each cell, and each line represents a single cell trace over time (n= 2-13 cells).

(E) Median intensity of mVenus in positive cells (determined by mVenus expression higher than the maximum mVenus intensity of the 0ng/mL doxycycline control) over time following addition of different concentrations of doxycycline (Dox). Median intensity levels per time point of mVenus are normalised to median intensity levels of the mScarleti nuclear marker, binned into 10 hour panels. N=2-3 independent experiments, n=8-90 per boxplot. Kruskal-wallis plus Dunn's posthoc test for multiple comparisons.

(F) The percentage of cells expressing mVenus following Tet-On activation with different concentrations of doxycycline (dox) per time point, binned into 10 hour panels. A positive cell is considered here any cell with a mean mVenus intensity above the maximum mVenus mean intensity for the 0ng/mL doxycycline sample for that particular experiment. N=2-3 independent experiments, n=2674-31234 cells over 30 timepoints per boxplot. Kruskal-wallis plus Dunn's posthoc test for multiple comparisons.

(G) Log likelihood ratio (LLR) of mVenus positive cells that pass as oscillators in Fig.1C at three different concentrations of doxycycline (Dox): 20 (n= 15), 200 (n=30), and 2000 (n= 117) ng/mL. N = 2-3 independent experiments. Kruskal wallis p=0.29 plus Dunn's posthoc test for multiple comparisons.

(H) Periodicity of mVenus oscillations in mVenus positive cells that pass as oscillators in Fig.1C at three different concentrations of doxycycline (Dox): 20 (n= 15), 200 (n=30), and 2000 (n= 117) ng/mL. N = 2-3 independent experiments. Kruskal wallis p=0.39 plus Dunn's posthoc test for multiple comparisons.

(I) Maximum fold change of the oscillations in mVenus positive cells that pass as oscillators in Fig.1C at three different concentrations of doxycycline (Dox): 20 (n= 15), 200 (n=30), and 2000 (n= 117) ng/mL. N = 2-3 independent experiments. Kruskal wallis p=0.0051 plus Dunn's posthoc test for multiple comparisons.

**Figure S2. Characterisation of dTAG dynamics and the ability to use dTAG to modify and create oscillatory protein expression. Related to Figure 2.**

(A) Mean of single cell traces showing mVenus intensity just before and following addition of different concentrations (above panels) of different dTAG drugs (right of panel). mVenus intensity is normalised to intensity before addition of the drug. The vertical dashed line indicates time at which drug was added. N = 2 independent experiments, n = 6 – 22.

(B) Density plot showing mScarlet intensity across a population of cells treated with different concentrations of 3 different dTAG drugs for a duration of 10 hours (30 timepoints). N = 2 independent experiments.

(C) Density plot showing mScarlet intensity across a population of cells after release from 3 different dTAG drugs for a duration of 10 hours (30 timepoints). N = 2 independent experiments.

(D) Single cells sequentially exposed to 2000 ng/mL doxycycline, 10nM dTAG$^{V}$-1 (blue shaded area), and re-addition of 2000 ng/mL doxycycline. N=25 cells.

(E)-(H) Single cell examples of PANC-1 UBC-NGN3-mVenus-FKBP-V cells exposed to periodic dTAG$^{V}$-1 addition. mVenus intensity is plotted, with the median background intensities removed. Cells were periodically exposed to media containing different concentrations of dTAG$^{V}$-1 (G-H; blue shaded areas), or DMSO as the control (E, grey shaded areas). Concentration of dTAG$^{V}$-1 is shown at the top of the plot (1nm (F), 10nm (G), or 1000nm (H)), and 6 cell examples shown from each concentration.

**Figure S3. Characterisation of the mathematical model, allowing us to predict optimal drug schedules for particular oscillatory behaviours. Related to Figure 3.**

(A) Model simulations (solid lines) of normalised NGN3 with optimised parameters at varying drug dosages, and drug-ON windows of [1 hr - 3 hrs] and [6.33 hrs - 12 hrs]. Mean normalised NGN3 Data from Fig.2H are in dashed lines.

(B) Plot of the Active degradation rate in equation (1) as a function of dTAG$^{V}$-1 drug dosage (taken from optimal parameter values found from A). This illustrates our modelling assumption that degradation caused by the addition of the drug does not scale linearly with dosage, but in a sigmoidal shape, and that degradation will be saturated at high nm concentrations. Note that the x-axis is plotted on a log-scale. Colours correspond to the drug concentrations shown in (A), blue = 1nm, green = 10nm and yellow = 1000nm dTAG$^{V}$-1.

(C) Schematic illustrating drug levels over time $D(t)$ as used within our mathematical model. Blue shaded areas indicate Drug-on windows, black line shows drug levels which are assumed to instantaneously reach intended dosage level $d$ at each $T^{ON}i$ time. Drug levels are then assumed to fall back to zero in accordance with first order decay kinetics upon each $T^{OFF}i$ time representing the washing out of drug from the well.

(D) Heatmap showing percentage difference between mean levels of model simulation and sinusoidal test data (with 5 hour period) at each drug-ON/drug-OFF time window.

(E) Heatmap showing percentage difference between the mean peak-to-trough fold-change (ratio between levels at peak and trough of wave) of model simulation and sinusoidal test data (with 5-hour period) at each drug-ON/drug-OFF time window.

(F) Heatmap showing percentage difference between the mean peak-to-peak period (time between subsequent peaks of wave) of model simulation and sinusoidal test data (with 5-hour period) at each drug-ON/drug-OFF time window.

(G) Scatter plot of drug-ON/drug-OFF times with marker positioning indicating the window schedule at which the model best fit to sinusoidal test data with periods 1-16 hrs. Colours of the markers indicates the sum of squares residual at these times. The dashed line shows symmetric drug-ON/drug-OFF times. This panel illustrates that, with fixed parameters, shorter drug-ON than drug-OFF times are required to mimic purely sinusoidal test data for shorter periods (e.g. 2 hrs). Conversely, longer drug-ON than drug-OFF times are needed to best fit the test data with longer periods (e.g. 16 hrs). Further, this shows that with fixed parameters, simulations are more perfectly sinusoidal at certain periodicities, e.g. 5hrs for our parameter set.

(H,J) Heatmaps showing sum of squares residual error between model simulation and sinusoidal test data with periods of 16 hrs and 2 hrs respectively across a grid of drug-ON/drug-OFF times. White circle indicates the point at which residual is minimised.

(I,K) Simulations of mathematical model (blue wave) determined at the drug-ON/drug-OFF time schedule which provided the best fit with sinusoidal test data at periods of 16 hrs and 2 hrs as shown by white circles in H and J.
